# Dissociating cognitive and motoric precursors of human self-initiated action

**DOI:** 10.1101/326686

**Authors:** N. Khalighinejad, E. Brann, A Dorgham, P. Haggard

## Abstract

Across-trial variability of EEG decreases more markedly prior to self-initiated than prior to externally-triggered actions, providing a novel neural precursor for volitional action. However, it remains unclear whether this neural convergence is an early, deliberative stage, or a late, execution-related stage in the chain of cognitive processes that transform intentions to actions. We report two experiments addressing these questions. Self-initiated actions were operationalized as endogenous ‘skip’ responses while waiting for target stimuli in a perceptual decision task. These self-initiated ‘skips’ were compared to blocks where participants were instructed to skip. EEG variability decreased more markedly prior to self-initiated compared to externally-triggered ‘skip’ actions, replicating previous findings.

Importantly, this EEG convergence was stronger at fronto-midline electrodes than at either the electrode contralateral or ipsilateral to the hand assigned to the ‘skip’ action in each block (Experiment 1). Further, convergence was stronger when availability of skip responses was ‘rationed’, encouraging deliberate planning before skipping (Experiment 2). This suggests that the initiation of voluntary actions involves a bilaterally-distributed, effector-independent process related to deliberation. A consistent process of volition is detectable during early, deliberative planning, and not only during late, execution-related time windows.

## Introduction

Our everyday actions span a spectrum of autonomy, ranging from simple reflexes, immediate motor responses to external stimuli, to complex volitional actions, which are not directly determined by any identifiable external stimulus (Haggard 2008). Functional and neuroanatomical studies have helped to refine this spectrum, identifying specific neural differences between self-initiated and externally triggered actions (Passingham 1987; Deiber et al. 1999; Jenkins et al. 2000; Cunnington et al. 2003; Passingham et al. 2010). An action that involves directly responding to an external cue, such as a choice reaction time, or responding to a verbal command, can be considered exogenous, or “externally-triggered”. In contrast, an endogenous, self-initiated action requires no external cue: the agent decides for themselves whether and when to act. Depending on the situation in which they are made, self-initiated actions often have additional distinctive properties linking them to voluntariness, and individual autonomy. For example, self-initiated actions are often susceptible to reason (Anscombe 2000), free from immediacy (Gold and Shadlen 2007), and often involve choosing between alternatives (Pereboom 2011).

The readiness potential (RP), also known as the bereitschaftspotential, is a gradual build-up of electrical potential in the pre-motor areas that occurs prior to self-initiated movements (Kornhuber, H. H., & Deecke 1965), often beginning one second or more prior to movement onset (Haggard 2008). Classically, RP is taken to be the electrophysiological sign of planning, preparation, and initiation of voluntary actions (Kornhuber, H. H., & Deecke 1990) and was suggested as the neural precursor to conscious intention to act (Libet et al. 1983).

The view that RP reflects a fixed precursor process that leads to self-initiated, voluntary action has been recently challenged (Schurger et al. 2012, 2016). These new models suggest that rising ramp pattern of the mean RP does not reflect a specific goal-directed process but rather reflect subthreshold fluctuations in premotor activity that influence the precise time of the action (Schurger et al. 2012; Murakami et al. 2014). The negative shape of the RP is not a readout of a specific preparatory process, but results from cross-trial averaging of these stochastic fluctuations time-locked to action-initiation, a common practice in computing mean RP.

Given the current debate on neural precursors of human voluntary action, we recently calculated the time-course of variability across trials in the RP signal. Classical models, which view the RP as marking a distinctive cognitive process preceding self-initiated actions, seem committed to low EEG variability during the RP period. In contrast, stochastic models would imply higher variability during the same period, with the proviso that variability must inevitably decrease prior to action, if the trigger for action indeed involves a neural signal approaching a threshold value.

Thus, the timing and magnitude of EEG variability prior to action can be informative about the neural processes that generate action. We indeed observed that across-trial variability of EEG decreased prior to action initiation. Further, this decrease was more marked prior to endogenous, self-initiated actions compared to externally-triggered actions (Khalighinejad et al. 2018). We showed that *in addition* to stochastic fluctuation in neural activity, a process of noise control prior to self-initiated actions may consistently contribute to decision time to move in humans (Khalighinejad et al. 2018). Importantly, this process could be observed as across-trial convergence of neural activity recorded by scalp electrodes from premotor cortex. However, it is not clear whether this neural marker of self-initiated action represents an effector-independent, general neurocognitive process of volition, or an effector-specific motoric activity.

RP begins symmetrically. However, prior to movement the RP lateralises, with stronger amplitudes observed over the hemisphere contralateral to the effector performing the movement (Eimer 1998). This lateralised readiness potential (LRP) reflects preparation to execute an action in an effector-specific manner (e.g. which hand will be used to press the button) (Kutas and Donchin 1980). In serial models of action control, LRP onset represents a useful dividing line between earlier ends and later means, separating the cognitive mechanisms of preparing actions, from the motor mechanisms related to the specific movement that implements the action goal. Here we ask whether the EEG convergence that precedes self-initiated action reflects an early cognitive, effector-independent process, or rather reflects a motoric, effector-dependent process.

To answer this question, we performed two experiments using a modified version of a recently-developed paradigm (Khalighinejad et al. 2018). In the original task, participants responded to the direction of unpredictably-occurring dot motion stimuli by pressing either the left or right arrow keys. Importantly, they could also choose to skip waiting for the stimuli to appear, by pressing both keys simultaneously whenever they wished. The skip response thus reflected a purely endogenous decision to act, without any direct external stimulus, and provided an operational definition of a self-initiated action. The self-initiated skip responses were compared to an externally-triggered skip block in which participants made the same skip action in response to an unpredictable visual cue, as opposed to whenever they wished. In the first experiment, we required participants to use either the left or right hand, in separate blocks, to perform the skip response. This modification enabled us to record EEG convergence contralateral to the acting hand. If our putative signal of self-initiated action is effector-specific, we would expect to see a stronger convergence at EEG electrodes contralateral to the hand assigned to perform the self-initiated skip response. Alternatively, if the signal is effector-independent, we would expect to see a stronger EEG convergence at midline electrodes, regardless of the hand used for skip response. This manipulation effectively probes the role of motoric execution-related processes in the initiation and elaboration of voluntary action.

In the second experiment, we ‘rationed’ volition, by restricting the number of self-initiated skip actions that could be made. We reasoned that this ‘rationing’ encourages careful and deliberate planning of self-initiated actions compared to a condition where ‘skip’ actions are unlimited. The logic was as before. If we find stronger convergence in the more deliberate, rationed condition, we would conclude that such convergence reflects cognitive processes that deliberate the likely value of action.

## Materials and methods

### Participants

The sample size calculation was obtained based on our previous experiment (Khalighinejad et al. 2018). 55 right-handed participants (23 for Exp.1 and 22 for Exp.2) aged 18-42 years old (17 male, mean age = 24.6 years) were recruited via the UCL Institute of Cognitive Neuroscience subject data pool. Six participants were excluded (three from each study), before data analysis, due to poor EEG quality. All subjects fulfilled the recruitment requirements, including no history of psychiatric and/or neurological disorders, no brain stimulation 48 hours prior to the study, having normal or corrected to normal vision, and no colour blindness. Participants received payment for their time based on an institution-approved hourly rate. The experimental design and procedure were approved by the UCL research ethics committee, and followed the principles of the Declaration of Helsinki.

### Behavioural task and procedure

Following consent, subjects sat in front of a computer screen (60 Hz refresh rate), in an electrically shielded chamber, and were fitted with the EEG cap. Participants were given verbal instructions and performed two practice blocks of trials to familiarise themselves with the task.

The behavioural task was adapted from our previous study (Khalighinejad et al. 2018). In brief, participants were required to focus on a fixation point in the centre of the screen. At the beginning of each trial the fixation point was black, but gradually and randomly changed colour. Meanwhile, participants observed randomly moving dots displayed within a circular aperture of 7o in diameter (density of 14.28 dots/degree, initially moving with 0% coherence with a speed of 2o/s) (Desantis, Waszak, and Gorea 2016; Desantis, Waszak, Moutsopoulou, et al. 2016).

Participants were instructed to wait until all dots moved in the same direction (step change to 100% coherence). The dots moved either to the left or right of the midline ranging in their degree of discrimination difficulty, with dot movement oriented upwards being more difficult to discriminate compared to sideways movements. The participant’s task was to determine the direction of dot motion, responding using the appropriate left (‘D’) or right (‘L’) keys on the keyboard with their respective left or right index finger. If participants responded correctly they were rewarded (2 pence), whilst incorrect responses resulted in a penalty (−1 pence). Responses that were early (before coherent dot motion) or delayed (after 2 s of coherent dot motion) also received a penalty (−1 pence) and were followed by an error message (Fig. 1).

**Figure 1.**
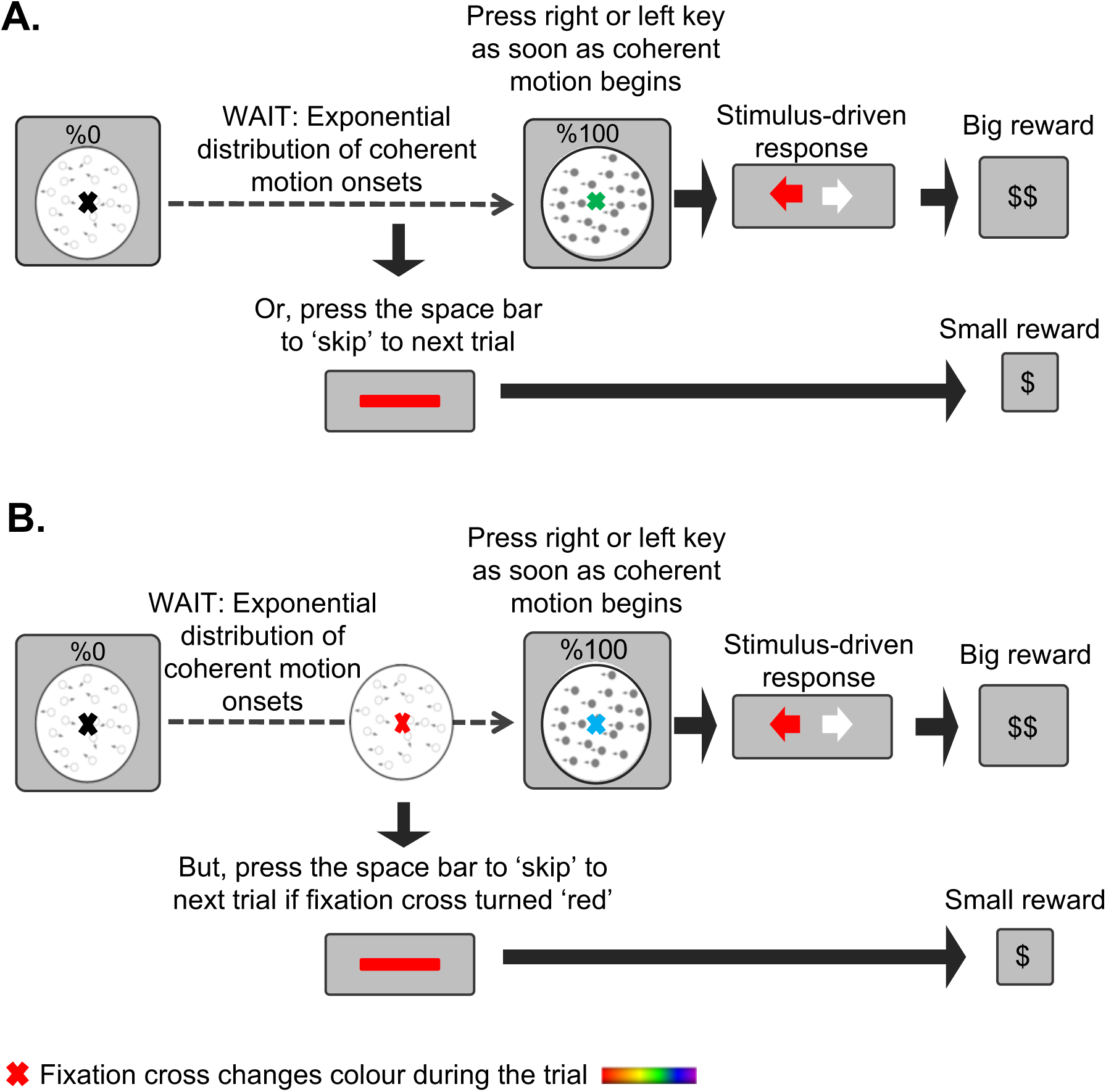
Timeline of an experimental trial. Participants responded to the direction of dot-motion with left and right keypresses. Dot-motion could begin unpredictably, after a delay drawn from an exponential distribution. A. In the ‘self-initiated’ blocks participants waited for an unpredictably occurring dot-motion stimulus, and were rewarded for correct left-right responses to motion direction. They could decide to skip long waits for the motion stimulus, by pressing the space bar with the left or right thumb (Experiment 1) or making a bilateral keypress (Experiment 2). They thus decided between waiting, which lost time but brought a large reward, and ‘skipping’, which saved time but brought smaller rewards. The colour of the fixation cross changed continuously during the trial, but was irrelevant to the decision task. B. In the ‘externally-triggered’ blocks, participants were instructed to press the space bar with left or right thumb (Experiment 1) or both (Experiment 2) when the fixation cross became red, and not otherwise.

Each experimental session was limited to 40 min. Participants were informed that the elapsing time between trial onset and coherent dot motion onset was random (drawn from an exponential distribution with min = 2 s, max = 60 s, mean = 12 s), and consequently the waiting time could be very long. To avoid long waiting times participants had the option to skip from one trial to the next by pressing the ‘space’ bar with their left or right thumb or both (Depending on the experiment. See later) and claiming a smaller reward (1 pence). They were reminded that they should consider the time-money trade off in each trial. They could win a big reward by waiting, if responded correctly, but loosing time, or could save time by skipping and collecting more guaranteed smaller rewards.

The task involved two alternating blocks that varied in skipping behaviour. In the *‘self-initiated’* blocks (Fig. 1A), participants could skip at any moment of their choosing following the onset of the trial. This skip response thus reflects a purely endogenous decision to act, in the absence of any external instruction to act, and based on the trade-off between later, larger, and smaller, earlier rewards. This provides an operational definition of self-initiated action within our experimental design. In the *‘externally-triggered’* blocks (Fig. 1B), participants had to skip in response to an external cue. The external cue was an unpredictable change in the colour of the fixation point to red. The colour cycle of the fixation cross had a random sequence and the timing of appearance of the red colour was yoked to the time of participant’s own previous skip responses in the immediately preceding self-initiated block. If participants skipped correctly in response to the colour cue they were rewarded (1 pence), however, they received a penalty (−1 pence) if they skipped too early (before the fixation point turned red) or too late (after 2 s of the fixation point being red). Thus, in *‘externally-triggered’* condition blocks, participants could not choose for themselves when to skip. To control for any confounding effect of attending to fixation point, participants were also required to monitor its colour in the self-initiated blocks and to roughly estimate the number of times the fixation point turned yellow, according to the following categories: never, less than 50%, 50%, more than 50%. The red colour was left out of the colour cycle in the self-initiated blocks.

At the end of each block, participants received feedback on their reward values, total elapsed time, and number of skips, enabling them to adjust their behaviour over time and maximise earnings. Each block consisted of 10 trials and the order of the blocks was counterbalanced across participants.

In Exp.1, participants used either the left or right thumb to perform the skip action. Once 40 min passed using one thumb, the task finished and subjects repeated the same task, after 5 min break, but using the other thumb to skip. Skipping hand order (right-left or left-right) was counterbalanced across participants. To ensure sufficient data collection and to avoid long testing sessions, the experiment was split across two sessions, which took place on different days. Data from similar conditions were pooled across the two sessions. Thus, the experiment had two factors: The hand used for skipping (left vs. right), and the type of skip action (self-initiated vs. externally-triggered). To perform the skip action in Exp.2, participants used both thumbs simultaneously. Exp. 2 consisted of two sessions: In the first session (*unlimited session*), similar to Exp.1, there was no limitation in the number of skip actions participants could perform in the self-initiated blocks. In the second session (*limited session*), however, they were informed that they could only make half the number of skips made in the first session. For example, if a participant skipped waiting 100 times in the first session, they were allowed to skip max 50 times in the second session. Total number of allowed skips was displayed on the screen at the beginning of the session and they were informed of the number of remaining skips at the end of each block. If a participant used all their allowed skip actions before the end of the experiment, the experiment continued but they were not allowed to skip anymore and had to wait for coherent dot motion before responding. They were not rewarded for saving their skip actions. The behavioural task was designed in Psychophysics Toolbox Version 3 (Brainard 1997).

### EEG recording

The experiment was conducted inside an electrically shielded chamber. EEG signals were recorded and amplified using an ActiveTwo Biosemi system (BioSemi, Amsterdam, The Netherlands). Participants wore a 64-channel EEG cap. To reduce preparation time, they were only fitted with a subset of 20 electrodes covering the central and visual areas: F3, Fz, F4, FC1, FCz, FC2, C3, C1, Cz, C2, C4, CP1, CPz, CP2, P3, Pz, P4, O1, Oz, O2. Horizontal and vertical electro-oculogram (EOG) recordings were made using external bipolar channels positioned on the outer canthi of each eye as well as superior and inferior to the right eye. Reference electrodes were positioned on the mastoid bone behind the right and left ears. EEG signals were recorded at a sampling rate of 2048 Hz. A trigger channel was used to mark the time of important events on the signal.

### EEG preprocessing

EEG data was preprocessed using Matlab (MathWorks, MA, USA) and EEGLAB toolbox (Delorme and Makeig 2004). Data was downsampled to 256 Hz and low-pass filtered at 30 Hz with no high-pass filtering. The average signal of the mastoid electrodes was used as a reference for all other electrodes. In Exp.1, EEG epochs for the hand used for skipping (left vs. right) and action conditions (self-initiated vs. externally-triggered) were separated resulting in four different types of epochs: self-initiated skip with left hand, self-initiated skip with right hand, externally-triggered skip with left hand, and externally-triggered skip with right hand. EEG epochs in Exp.2 were separated based on the action condition (self-initiated vs. externally-triggered) and the availability of skip actions (unlimited vs. limited). Each epoch was 4 s long, ranging from 3 s prior to 1 s after skip action. Overlapping epochs, in which participants skipped earlier than 3 s from trial initiation were removed. Next, non-ocular artefacts were removed from data using two methods of improbable data rejection (single channel SD = 6, global SD = 2) and extreme values rejection (upper threshold limit = 250 µV, lower threshold limit = -250 µV). Following this, ocular artefacts components were extracted by independent component analysis (ICA) and were identified and removed by visual inspection. Trials with artefacts remaining after this procedure were excluded by visual inspection. In keeping with our previous study (Khalighinejad et al. 2018), RP recordings were baseline corrected using a baseline of -5 to +5 ms with respect to action onset, avoiding any assumptions concerning the moment of RP initiation.

### EEG analysis

EEG analysis was conducted using Matlab (MathWorks) and FieldTrip toolbox (Oostenveld et al. 2011). Mean RP amplitude across trials and variability of RP amplitudes across trials (measured by SD) were measured as dependent variables for each trial type. We previously showed that in a bimanual task EEG convergence is strongest in fronto-midline electrodes (Khalighinejad et al. 2018). However, this topography might reflect either a midline cognitive process of deliberation, or the summation of effector-related motoric activity linked to controlling both hands simultaneously. Our current design, based on blocked, unimanual skip responses, was designed to distinguish between these possibilities. The FCz electrode was chosen as a marker of effector-independent cognitive processing, while C3 and C4 electrodes, positioned above the left and right motor cortex, respectively, were chosen as marking effector-dependent processes in the contralateral motor cortices. First, cluster-based permutation tests were used to compare across-trial SD between self-initiated and externally-triggered conditions, for each chosen electrode. The cluster-based permutation tests were performed using the following parameters: time interval = [-2 - 0 s relative to skip action], number of draws from the permutation distribution = 1000.

In a subsequent step, we investigated whether EEG convergence is an effector-dependent or -independent process. EEG convergence was defined as the area between the SD curve in self-initiated and externally-triggered condition in a two second window prior to skip action onset. EEG convergence from C3 with left hand skips was averaged with EEG convergence from C4 with right hand skips to give the ipsilateral EEG convergence. Analogously, EEG convergence from C3 with right hand skips was averaged with EEG convergence from C4 with left hand skips to give the contralateral EEG convergence. EEG convergence from FCz with left hand skips was averaged with EEG convergence from FCz with right hand skips to give frontal-midline EEG convergence. As we planned to compare these neurocognitive processes between electrodes selected on the basis of previous studies, we used one-way repeated-measures ANOVA.

In the second experiment, we did not have a priori hypotheses about where in the brain ‘rationing’ would be found, so we used a more conservative, exploratory approach. Briefly, non-parametric permutation tests across all electrodes from central areas (see *EEG recording*) were used to compare convergence between conditions in the second experiment. This approach avoids some of the arbitrary assumptions associated with electrode and time-bin selection. The permutation tests were performed using the following parameters: time interval = [-2-0 s], minimum number of neighbouring electrodes required = 2, number of draws from the permutation distribution = 1000.

## Results

### Experiment 1: EEG convergence is an effector-independent process

On average, participants (n=20) skipped waiting for coherent dot motion 63 times (SD = 3) with the left hand and 62 times (SD = 3) with the right hand in the self-initiated condition. They skipped 62 (SD = 3) and 63 (SD = 3) times with the left and right hand, respectively, in response to the external-cue. The mean waiting time before skipping was 6.68 s (SD = 1.04) for the left hand and 6.91 s (SD = 0.58) for the right hand in the self-initiated condition (Fig. 2.A,B). This waiting time in the externally-triggered condition was 7.22 s (SD = 1.11) and 7.46 (SD = 0.61), for the left and right hand, respectively. This confirms that our yoking procedure (see materials and methods) was successful. For externally-triggered trials, the reaction time to external cue (red fixation point) was 752 ms (SD = 76) and 742 ms (SD = 89) for left and right hand, respectively. Participants gained on average an additional 125p (SD = 2.77) for skipping with the left hand and 125p (SD = 4.01) for skipping with the right hand. They further gained 125p (SD = 48.90) and 127p (SD = 42.34) with left and right hand, respectively, from correct responses to coherent dot motion. No significant difference was observed in any of the above behavioural measures between the left and right hand (*p* > 0.27 for all comparisons) (see supplementary table 1 for full waiting time data).

**Figure 2.**
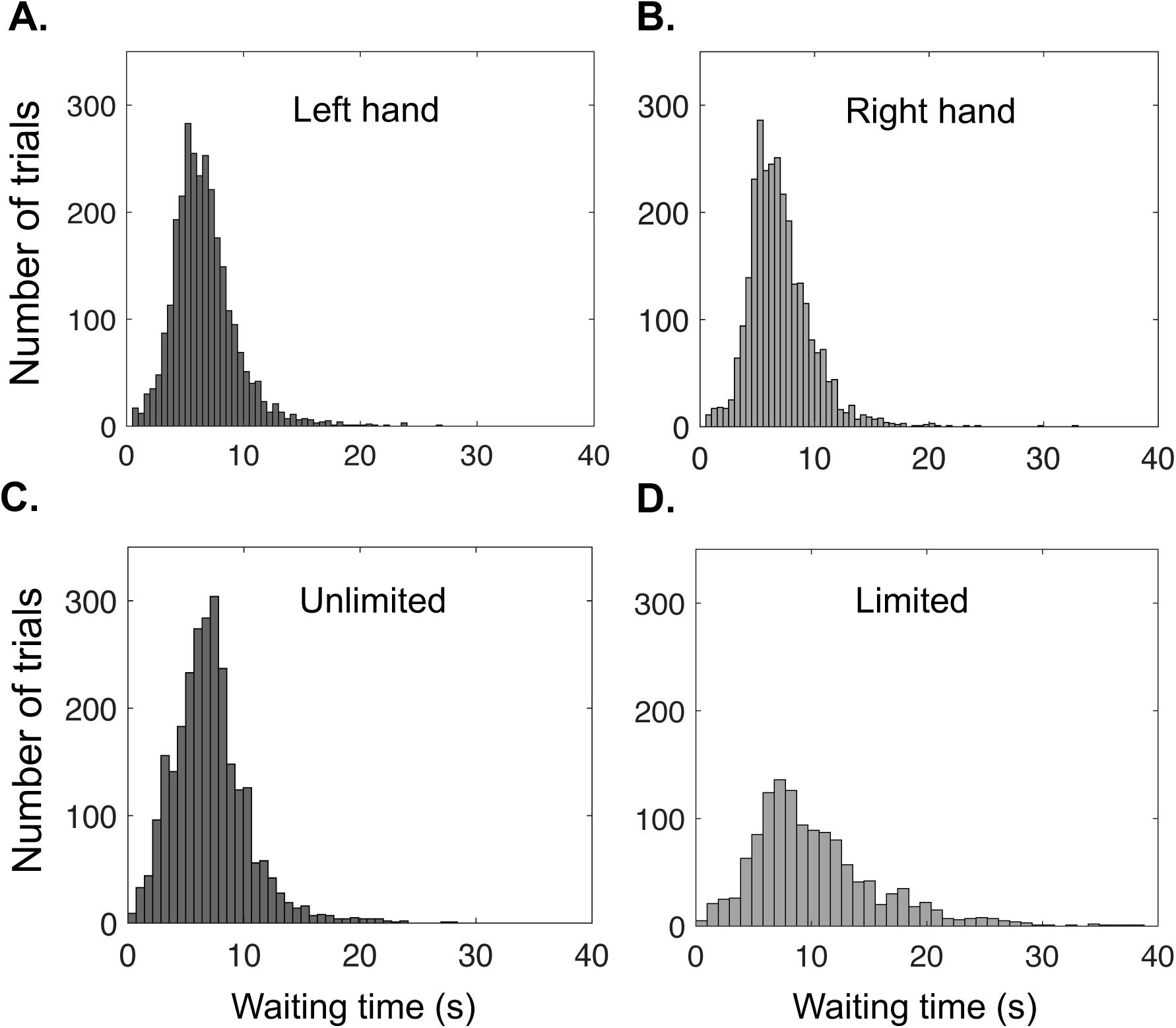
Histogram of waiting time before skipping in self-initiated condition. In Experiment 1 participants could skip waiting by pressing the space bar with their left (A) or right (B) hand. In Experiment 2 participants used their both hands simultaneously to skip waiting but in the ‘limited’ condition (D) they were only allowed to make half the number of skips they made in the ‘unlimited’ condition (C).

We next investigated whether reaction time to coherent dot motion was influenced by the hand assigned to skip response (e.g., how preparing to skip with the left hand influences the reaction time to rightward moving dots). We found that reaction time to dot motion direction was marginally lower when the hand used to respond to dot motion direction corresponded with the hand assigned for the skip response on that block (F (1,19) = 4.25, p = 0.05) (supplementary Fig. 1). This suggests that action initiation in response to an external cue (coherent dot motion) may be facilitated when precursor processes have already prepared for a self-initiated action (skip response).

EEG data from 20 participants was preprocessed and pooled across the two sessions. Fig.3.A,B and Fig.4.A,B,E,F show the grand average RP for each trial type, from the selected electrodes. The RP grand average showed the expected negative-going shape for self-initiated action for both left and right hand skip responses and at all electrodes of a priori interest. As expected, RP was more prominent in electrodes contralateral to the skipping hand (C3 for right hand skips (Fig. 4B) and C4 for left hand (Fig.4E)).

**Figure 3.**
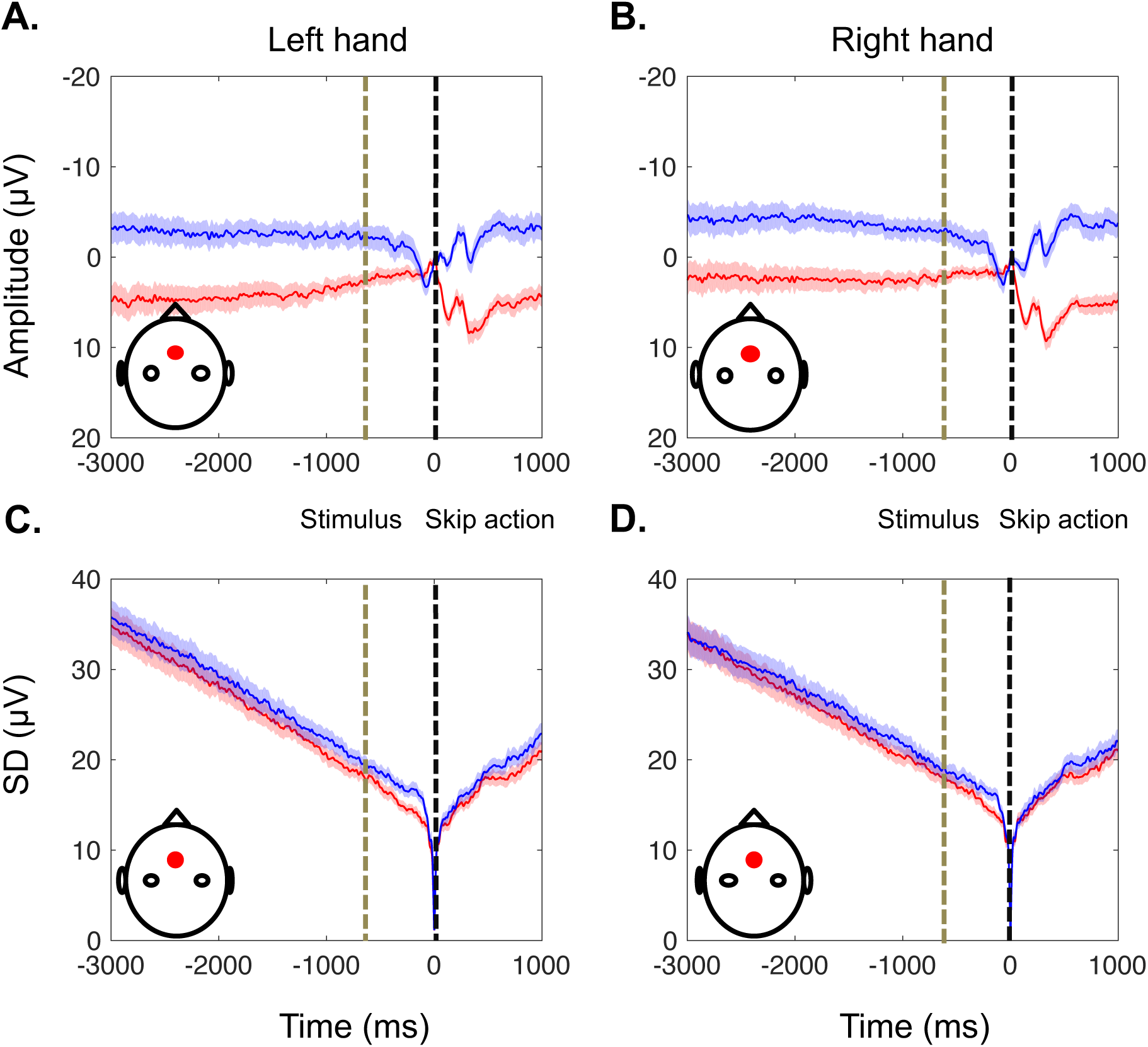
EEG activity from the effector-independent electrode (FCz) prior to skip actions. The red and blue lines represent self-initiated and externally-triggered skip conditions, respectively. Data is time-locked to the skip action (black vertical line), baseline-corrected in a 10 ms window around the skip. The average time of the skip instruction (fixation cross changing to red) in the externally-triggered condition is shown as a grey vertical line. A,B. Grand average RP amplitude ± standard error of the mean across participants (SEM) for the skip responses with the left (A) and right hand. C,D. Standard deviation across trials averaged across participants ± SEM for the skip responses with the left (C) and right (D) hand.

**Figure 4.**
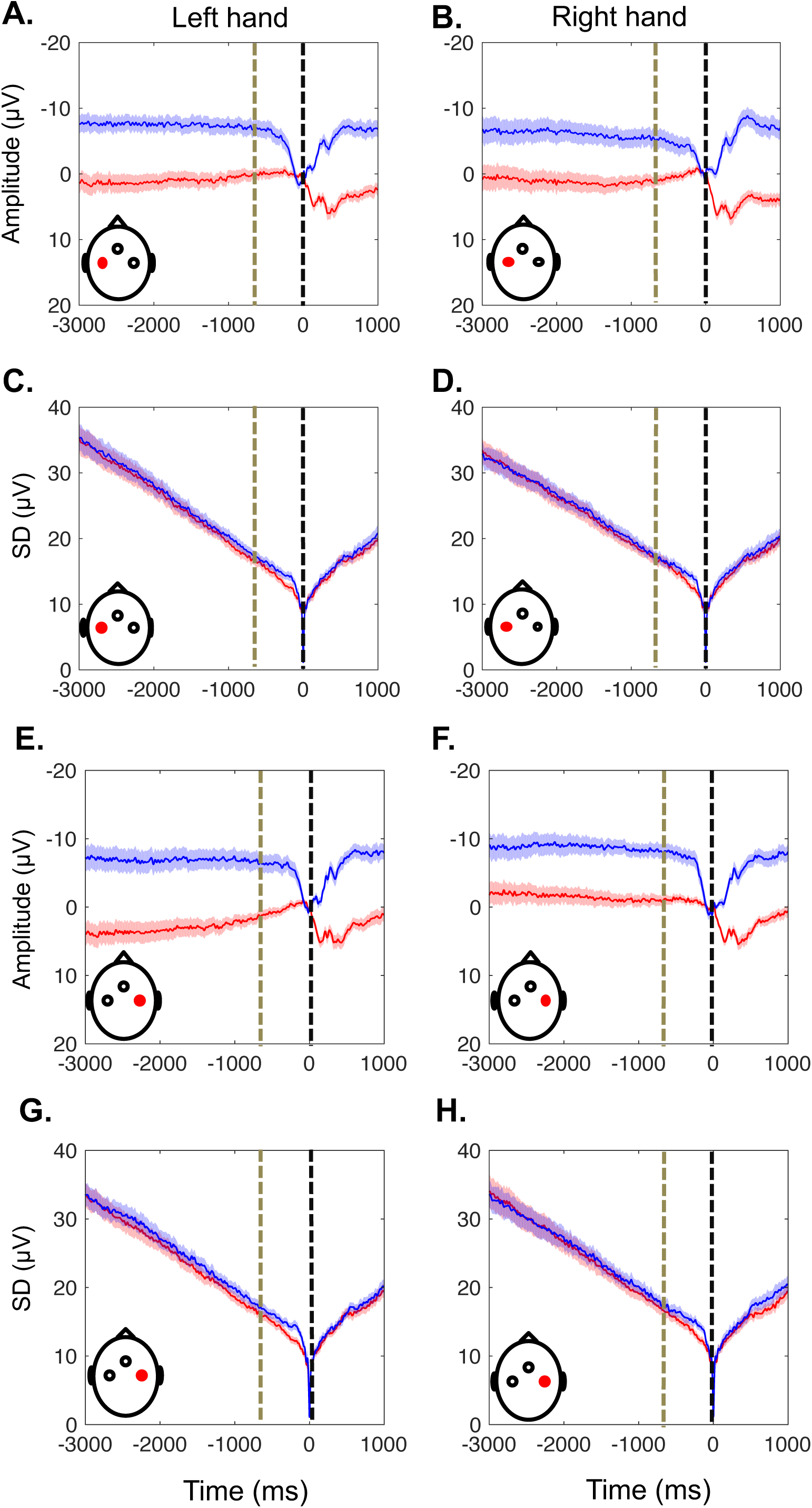
EEG activity from the effector-specific electrodes prior to skip actions. The red and blue lines represent self-initiated and externally-triggered skip conditions, respectively. Format as in Fig. 3. A,B. Grand average RP amplitude ± standard error of the mean across participants (SEM) from electrode C3 for the skip responses with the left (A) and right (B) hand. C,D. Standard deviation across trials averaged across participants ± SEM for the skip responses from electrode C3 with the left (C) and right (D) hand. E,F. Grand average RP amplitude ± standard error of the mean across participants (SEM) from electrode C4 for the skip responses with the left (E) and right (F) hand. G,H. Standard deviation across trials averaged across participants ± SEM for the skip responses from electrode C3 with the left (G) and right (H) hand.

**Figure 5.**
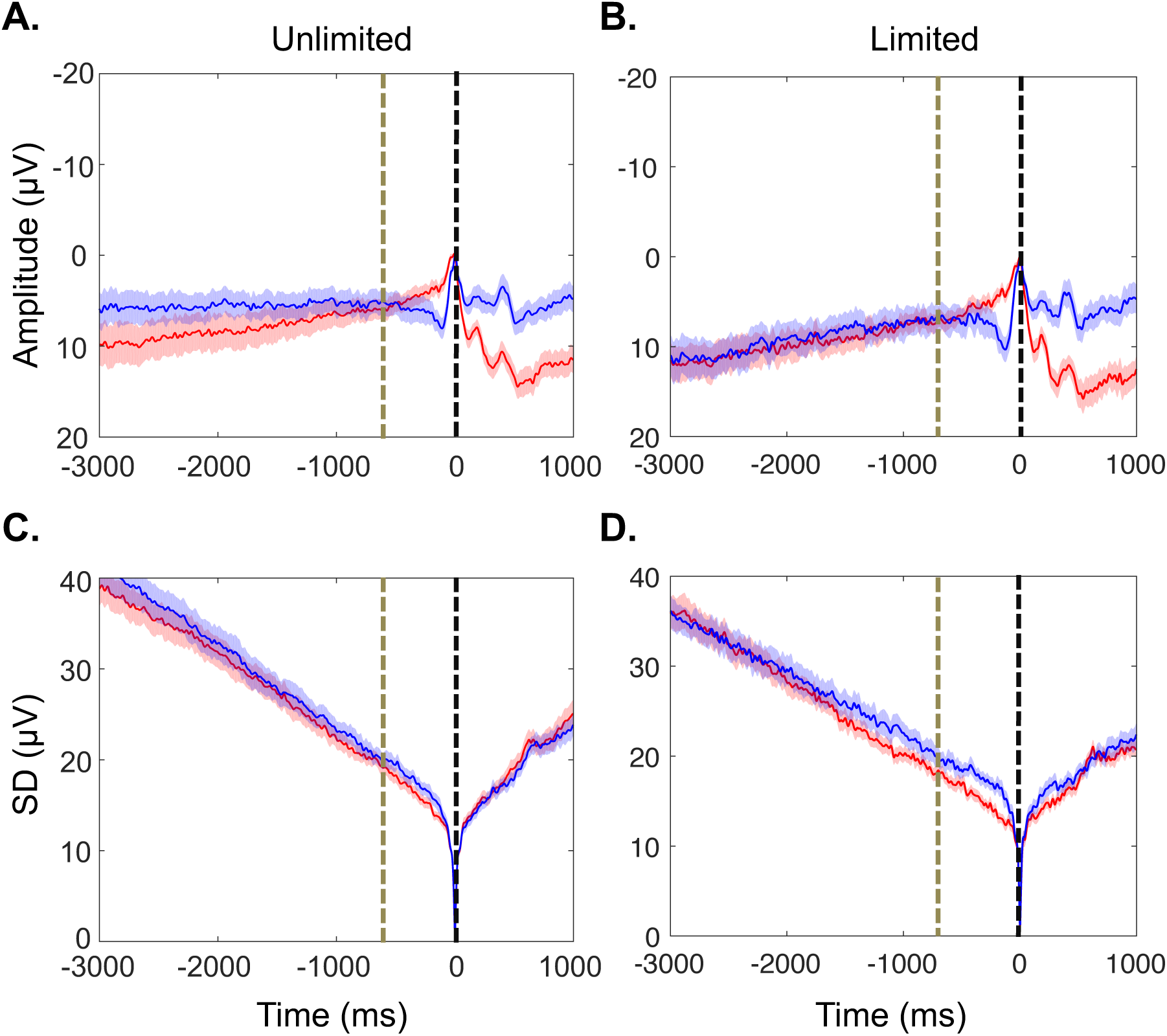
EEG activity prior to skip actions. The red and blue lines represent self-initiated and externally-triggered skip conditions, respectively. Data is time-locked to the skip action (black vertical line), baseline-corrected in a 10 ms window around the skip, and recorded from FCz electrode. The average time of the skip instruction (fixation cross changing to red) in the externally-triggered condition is shown as a grey vertical line. A,B. Grand average RP amplitude ± standard error of the mean across participants (SEM) for the Unlimited (A) and Limited (B) skip conditions. C,D. Standard deviation across trials averaged across participants ± SEM for the skip responses in the Unlimited (C) and Limited (D) skip conditions.

We first aimed to replicate our pervious results (Khalighinejad et al. 2018) and to validate the use of EEG convergence as a reliable marker of self-initiated action. Thus, inter-trial variability of RP amplitudes was calculated for each trial type (Fig. 3.C,D and Fig.4.C,D,G,H). The decrease in SD is partly an artefact of time-locking and baseline correction. However, this artefact is common to the self-initiated and externally-triggered conditions. Thus, the difference in convergence between the self-initiated and externally-triggered conditions suggests a precursor process that precedes self-initiated action (see Khalighinejad et al. 2018 for details). Cluster-based permutation testing was used to investigate whether SD decrease prior to self-initiated skip action is significantly different from SD decrease prior to externally-triggered skip action in our electrodes, in the latency range from -2s to the time of skip action onset (see materials and methods). Data from FCz (effector-independent) showed significant EEG convergence for both left (p < 0.02) and right hand (p < 0.01) skip responses. The same pattern was observed for effector-dependent electrodes. Specifically, the convergence was significantly different for left (p < 0.02) and right hand (p < 0.01) data from C3 and left (p < 0.02) and right hand (p < 0.05) data from C4. These findings confirm previous reports that neural activity prior to self-initiated actions gradually converges towards an increasingly stable pattern, and thus could be used as a reliable marker of self-initiated action.

To ensure that the key cognitive factors in the task were balanced between self-initiated and externally-triggered conditions, we also analysed the mean and SD of EEG amplitude prior to stimulus-triggered responses to coherent dot motion (as opposed to skip responses), for each responding hand and each electrode. We did not observe any negative-going potential or differential EEG convergence prior to coherent dot motion (p > 0.05) (see supplementary figures 2-4). This suggests that the disproportionate drop in SD prior to skip actions cannot be explained merely by a difference in background EEG, such as expectation of dot stimuli or temporal processing. That is, only those processes related to self-initiated action showed the distinctive EEG convergence, while general features of the context or block that were unrelated to the action event itself, did not show any altered EEG variability.

After replicating our previous results we aimed to investigate whether EEG convergence reflects an effector-independent, cognitive process, or an effector-specific motoric process. Based on the hand used for skipping, EEG convergence data (area between the SD curve in self-initiated and externally-triggered condition) was divided into three categories: *ipsilateral*, *contralateral* and *midline* EEG convergence (see materials and methods for definitions). A one-way repeated measures ANOVA showed a significant difference between these three categories (F(2,38) = 3.87, p = 0.030). The omnibus ANOVA was followed up by specific pairwise comparisons (no correction for multiple comparisons is required for this situation, following Fisher’s LSD procedure (Meier 2006)). Specifically, *midline* EEG convergence was stronger compared to both *ipsilateral* (t(19) = -2.60, p = 0.017) and *contralateral* (t(19) = -2.12, p = 0.048) EEG convergence. In contrast, there was no difference between the contralateral and ipsilateral EEG convergence (t(19) = 0.24, p = 0.816). These results suggest that the primary neurocognitive process measured by EEG convergence is not lateralised and does not depend on the specific effector used to perform a self-initiated action.

### Experiment 2: Deliberation about self-initiated action boosts neural precursor processes

If, as suggested by Experiment 1, our putative signal of self-initiated action is an effector-independent cognitive process, we would expect a stronger signal when this process is highlighted by key cognitive conditions that affect volition. In this experiment, we ‘rationed’ volition, by restricting the number of self-initiated actions that participants could use to skip long foreperiods. In the ‘unlimited’ condition participants could skip as many times as they wanted. On average they skipped 123 times (SD = 48). For the ‘limited’ condition they were instructed that they could make exactly half the number of skips they had previously made in the ‘unlimited’ condition. On average they made 59 skips (SD = 23). On average, participants waited 7.6 s (SD = 1.7) before skipping in the self-initiated ‘unlimited’ condition and 8.1 s (SD = 1.8) in the externally-triggered ‘unlimited’ condition. For the ‘limited’ condition they waited 11.3 s (SD = 2.2) and 11.8 (SD = 2.2) in the self-initiated and externally-triggered action blocks, respectively. Waiting time before skipping was significantly longer and had a wider distribution in self-initiated ‘limited’ compared to ‘unlimited’ condition (t(18) = 10.33, p < 0.01) (Fig. 2C,D). This suggests that rationing the number of *skip* actions encouraged careful planning of self-initiated actions, and a deliberate decision to use skip actions only where they were most valuable – i.e., when the wait for dot motion onset was particularly prolonged.

EEG data from 19 participants was preprocessed. The number of skip responses in the ‘limited’ condition was restricted by instruction to half the number of skip responses that the participant had previously made in the ‘unlimited’ condition. To control for this difference when measuring inter-trial variability, a subset of trials, equal to the number of skip responses of each participant in the ‘limited’ condition, was randomly extracted from the unlimited condition. All subsequent EEG analysis of ‘unlimited’ condition was performed on these subsets of trials. Fig.5.A,B shows the grand average RP for each trial type. The RP grand average showed the expected negative-going shape for self-initiated action in both ‘limited’ and ‘unlimited’ conditions. The difference in EEG convergence between the self-initiated and externally-triggered actions was significant in the ‘limited’ condition (p < 0.02, cluster-based permutation test across central electrodes) but not in the ‘unlimited’ condition (p > 0.05, cluster-based permutation test across central electrodes). Importantly, cluster statistics showed that this decrease in trial-to-trial variability was significantly more marked when the number of skip actions was limited compared to ‘unlimited’ condition (p < 0.05, cluster-based permutation test across central electrodes). This suggests that the putative precursor signal of self-initiated action is intensified when participants are encouraged to plan and deliberate ‘*when’* to initiate a volitional action.

## Discussion

We measured the convergence of individual trial EEGs towards a stable RP-like pattern prior to self-initiated ‘skip’ actions, and prior to unpredictable external signals instructing a skip. We used the degree of convergence as a proxy for the neural processes underlying self-initiated, voluntary action initiation. We found that this convergence was independent of the hand designated to execute the motor action, and predominantly involved frontal midline structures (experiment 1), suggesting it reflects cognitive preparation for volitional action, rather than motoric preparation for action execution. We also found that restricting the availability of self-initiated actions, which presumably lead to enhanced deliberation prior to action initiation due to greater action value, boosted this convergence (experiment 2). We conclude that EEG convergence prior to self-initiated action reflects a precursor neural process that involves deliberation and motivation for action, and not merely the motoric means of executing the action.

The common interpretation of the readiness potential as a causal signal for voluntary action has been recently challenged. New models suggest that the decision of ‘when to move’ in a self-initiated task is determined by stochastic fluctuation in neural activity (Schurger et al. 2012). Extending these new interpretations, we recently showed that self-initiated actions are preceded by a gradual convergence in neural activity towards an increasingly stable pattern. Interestingly, this convergence could be modelled within a stochastic fluctuation framework, but only by assuming an additional process of neural noise regulation (Khalighinejad et al. 2018). In the present study, by performing two separate experiments, we asked if convergence reflects a general, effector-independent, cognitive process, or an effector-dependent, lateralised motoric process. Experiment 1 aimed to investigate the origin in the brain of this convergence prior to self-initiated action. EEG variability decreased more markedly prior to self-initiated compared to externally-triggered actions, replicating our previous findings (Khalighinejad et al. 2018). This convergence in EEG was observed at electrodes over the motor cortices, and also at more frontal midline electrodes classically associated with RP. Importantly, control analyses ruled out the possibility that EEG convergence was merely some unknown contextual difference between the self-initiated and externally-triggered action conditions. Coherent dot motion starts at a random, unpredictable time. Therefore, any general contextual difference, not related to action preparation, should be captured by EEG variability time-locked to responses to coherent dot motion – yet no such differences were found (Supplementary figures 2-4). This suggests that reduced variability in self-initiated skip conditions is linked to the impending action itself, and not to any general difference in expectancy or task demands between the two conditions.

EEG convergence was stronger over effector-independent, midline electrodes (FCz electrode) compared to electrodes contralateral and ipsilateral to the hand used for skipping (C3 and C4 electrodes). This suggests that EEG convergence is frontally and bilaterally distributed. In particular, it arises upstream of the LRP, which has generally been localised to the primary motor cortex and lateral premotor cortex (Eimer 1998; Haggard and Eimer 1999). Thus, this precursor process of self-initiated action appears to be independent of how the action is motorically expressed. Our results therefore distinguish an early cognitive decision to make a self-initiated action, from the later motoric computations regarding *how* to implement and execute that action. Our findings favour a serial and hierarchical evolution of self-initiated action, with a specifically cognitive stage of deliberation, that occurs in advance of execution-related stages. This contrasts with other models, in which interactive competitive inhibition between motor execution plans might, in itself, constitute the decision process (Cisek 2012).

Recent models view action preparation and execution as two independent processes with distinct neural basis (Haith et al. 2016). However, self-initiated action execution and preparation may not necessarily be serial processes that follow each other in a deterministic way, but could rather be two independent overlapping processes, with the gradual flow of activity from the preparatory to execution state, as opposed to a sharp transition. For example, Orban de Xivry et al. (2016), measured reaction time during a virtual line bisection task in which participants used a robotic manipulandum to move a cursor smoothly through the middle of a target line that varied in its location and orientation. Movements to targets farther away from the starting position had shorter reaction times than movements to closer targets. They also observed that a larger increase in movement curvature from the nearer to farther target was associated with a larger reduction in reaction time. They concluded that, if processed before movement onset, the cognitive demands of accurately planning the movement gave rise to an increase in reaction time. However, delaying accuracy planning during execution led to a substantial decrease in reaction time. They thus argued that preparation for voluntary action might overlap in time with movement execution, although the two processes may still be informationally independent. For instance, an overlap period between preparation and execution may explain why one can still veto an internally-generated action even after the onset of action preparation (Schultze-Kraft et al. 2015).

Recently, Bozzacchi et al. (2016), studied the motor preparation of the two hands’ movements in a task where participants had to reach congruent or incongruent targets. Motor related cortical potentials showed that movements of both hands were programmed as a single motor plan, although they differentiated during movement execution depending on the target. A lack of lateralization in neural activity prior to movement onset, including in incongruent movement conditions in which each hand moved towards a different target, suggested that the two hands’ actions were planned as a whole, and not as separate movements. Interestingly, these authors used a self-paced task, where participants could freely choose when to initiate the movement. Thus, suggesting a possible link between the ‘single motor program’ in their study and the ‘effector-independent precursor processes’ that we identified.

While experiment 1 investigated the role of late, motoric processes, experiment 2 focussed instead on early, motivational processes for initiating voluntary action. We reasoned that, if our putative signal of self-initiated action indeed represents an early process linked to the motivation and value of initiating action, we might be able to boost this signal by encouraging participants to plan and deliberate over when to act. Participants waited longer, and EEG convergence was stronger when they could only perform a limited number of skip actions compared to a condition with no such limit. This effect could be driven by both an exaggerated EEG convergence in the ‘limited’ condition, as self-initiated actions become more deliberate and goal-directed, or by a diminished EEG convergence in the ‘unlimited’ condition, as self-initiated actions become more frequent, more routine, and more habitual. Our design and analysis cannot directly tease apart the opposing effects of deliberative volition on the one hand, and of habitual action on the other. However, in our ‘unlimited’ condition, reduction in inter-trial variability of self-initiated actions was not significantly different from that for externally-triggered actions. This may suggest that self-initiated actions became more habitual as they became more frequent, and then showed no more convergence or stability in the EEG pattern than control trials without self-initiated action. On this view, the distinctive pattern of neurocognitive activity prior to self-initiated action would reflect a specific process of deliberating whether or not to act now.

Interestingly, a recent study compared neural precursors of action for arbitrary and deliberate decisions (Maoz et al. 2018). While RPs were found for arbitrary decisions, they were absent in deliberate decisions. At first this may seem contrary to our findings. However, in that study the mean readiness potential was used as the precursor signal of action. We likewise found no difference between RPs in deliberate decisions of ‘limited’ condition and habitual decisions of ‘unlimited’ condition. But when using EEG convergence, rather than RP, as our putative signal of self-initiated action we found evidence for stronger processing prior to deliberate action decisions. This finding suggests that EEG convergence reflects a general cognitive process that could be intensified by careful planning and deliberation.

In summary, we showed that EEG convergence is a robust and reliable marker of self-initiated voluntary action. Importantly, this convergence in neural activity is a neurocognitive process that is bilaterally distributed and is effector-independent. The neural processes of self-initiated action appear to be independent of how the action is motorically expressed, but are sensitive to the reasons and motivation for initiating action. Cognitive models of intentionality propose a serial and hierarchical progression from prior intention to intention in action (Searle 2008), or from distal intention to proximal and motor intention (Pacherie 2008). Our finding shows that a consistent neural activity is associated with the earlier stages of this chain, and not only with final motor execution.

## Acknowledgments

This work was supported by the European Research Council Advanced Grant HUMVOL (Grant number: 323943), and by The Leverhulme Trust (Ref. RPG-2016-378). The authors declare no competing financial interest.

## Supplemental data

**Supplementary Figure 1.**
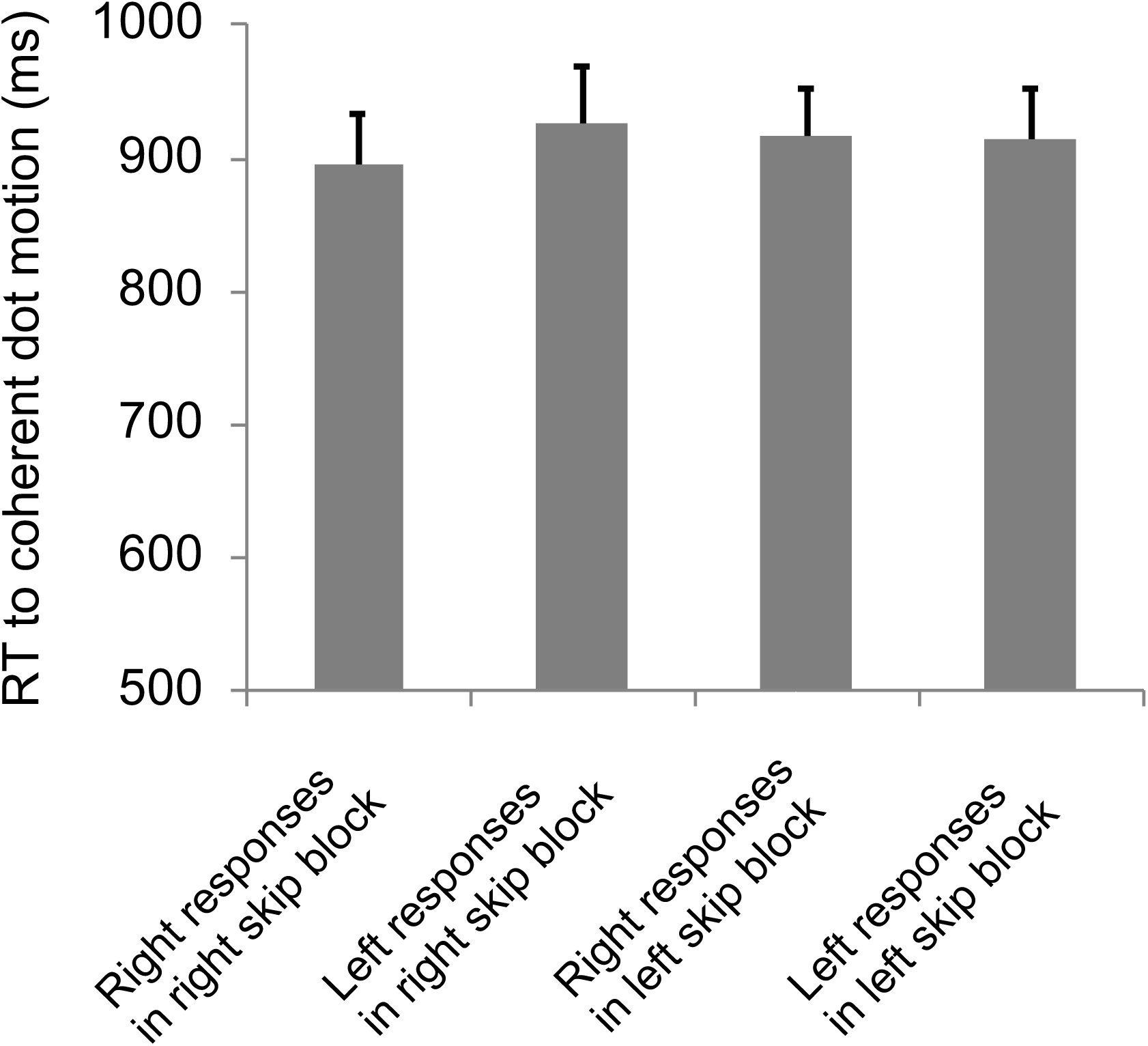
Reaction time to coherent dot motion onset. RT was shorter when the hand assigned to perform the skip response was the same as the direction of coherent dot motion (Right responses in right skip blocks, and left responses in left skip blocks). RT was longer when the hand assigned to perform the skip response was different from the direction of coherent dot motion (Right responses in left skip blocks, and left responses in right skip blocks).

**Supplementary Figure 2.**
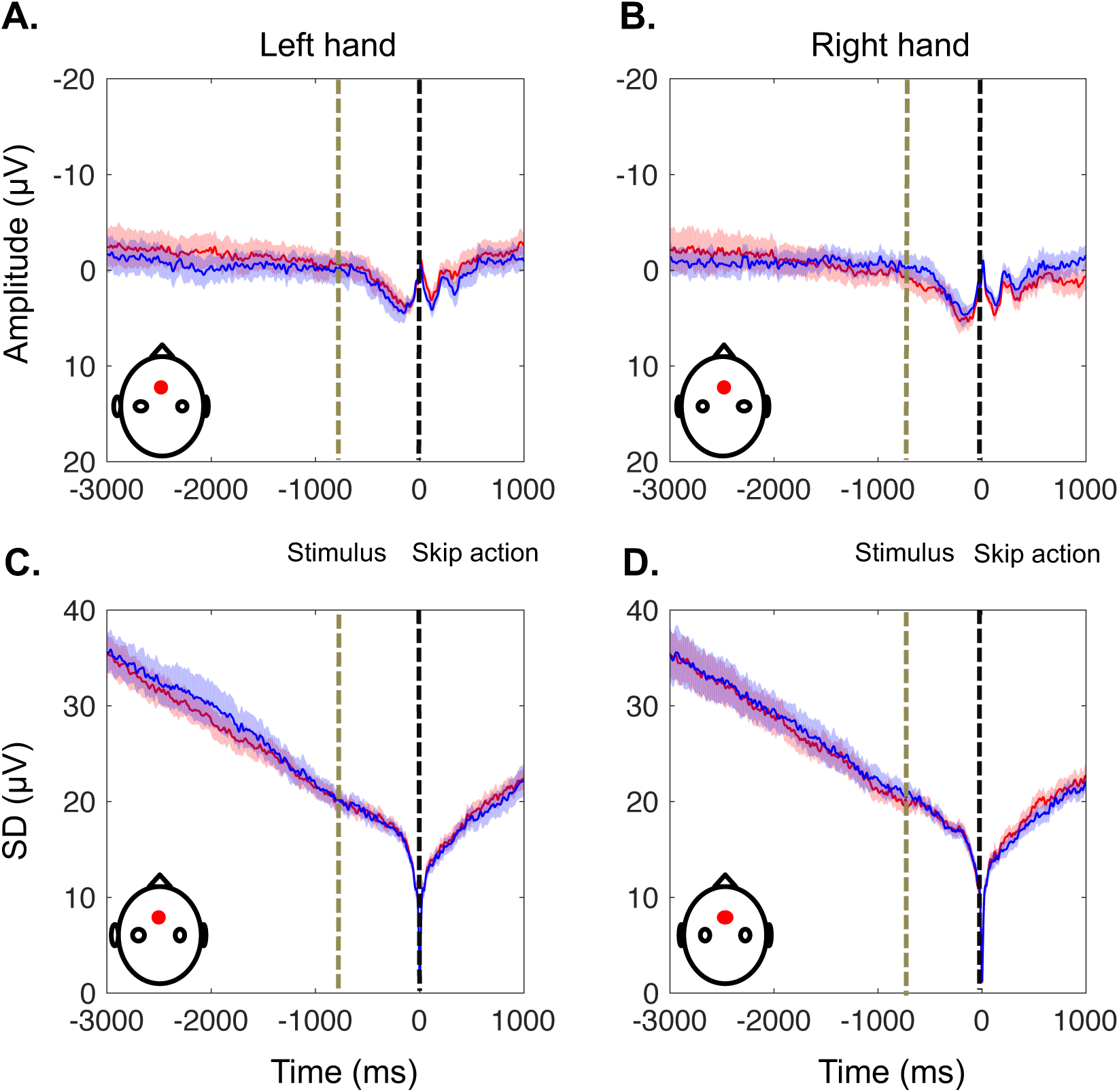
EEG activity from the effector-independent electrode (FCz) prior to response to coherent dot motion. The red and blue lines represent self-initiated and externally-triggered skip conditions, respectively. Data is time-locked to the response to coherent dot motion direction (black vertical line), baseline-corrected in a 10 ms window around the response. The average time of the coherent dot motion onset is shown as a grey vertical line. A,B. Grand average RP amplitude ± standard error of the mean across participants (SEM) for the responses with the left (A) and right (B) hand. C,D. Standard deviation across trials averaged across participants ± SEM for the responses with the left (C) and right (D) hand.

**Supplementary Figure 3.**
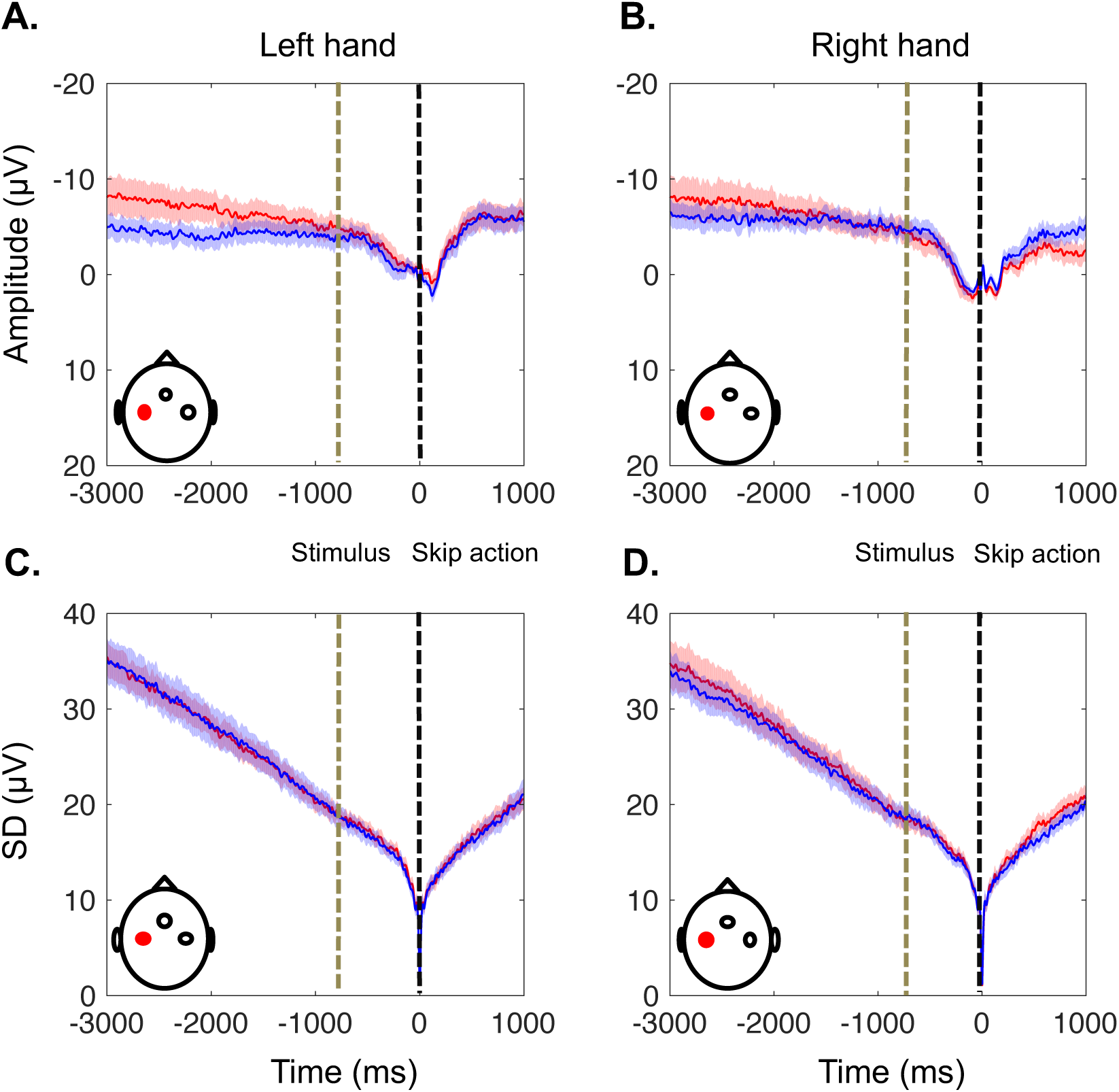
EEG activity from the effector-independent electrode (C3) prior to response to coherent dot motion. The red and blue lines represent self-initiated and externally-triggered skip conditions, respectively. Format as in Supplementary Fig. 2. A,B. Grand average RP amplitude ± standard error of the mean across participants (SEM) for the responses with the left (A) and right (B) hand. C,D. Standard deviation across trials averaged across participants ± SEM for the responses with the left (C) and right (D) hand.

**Supplementary Figure 4.**
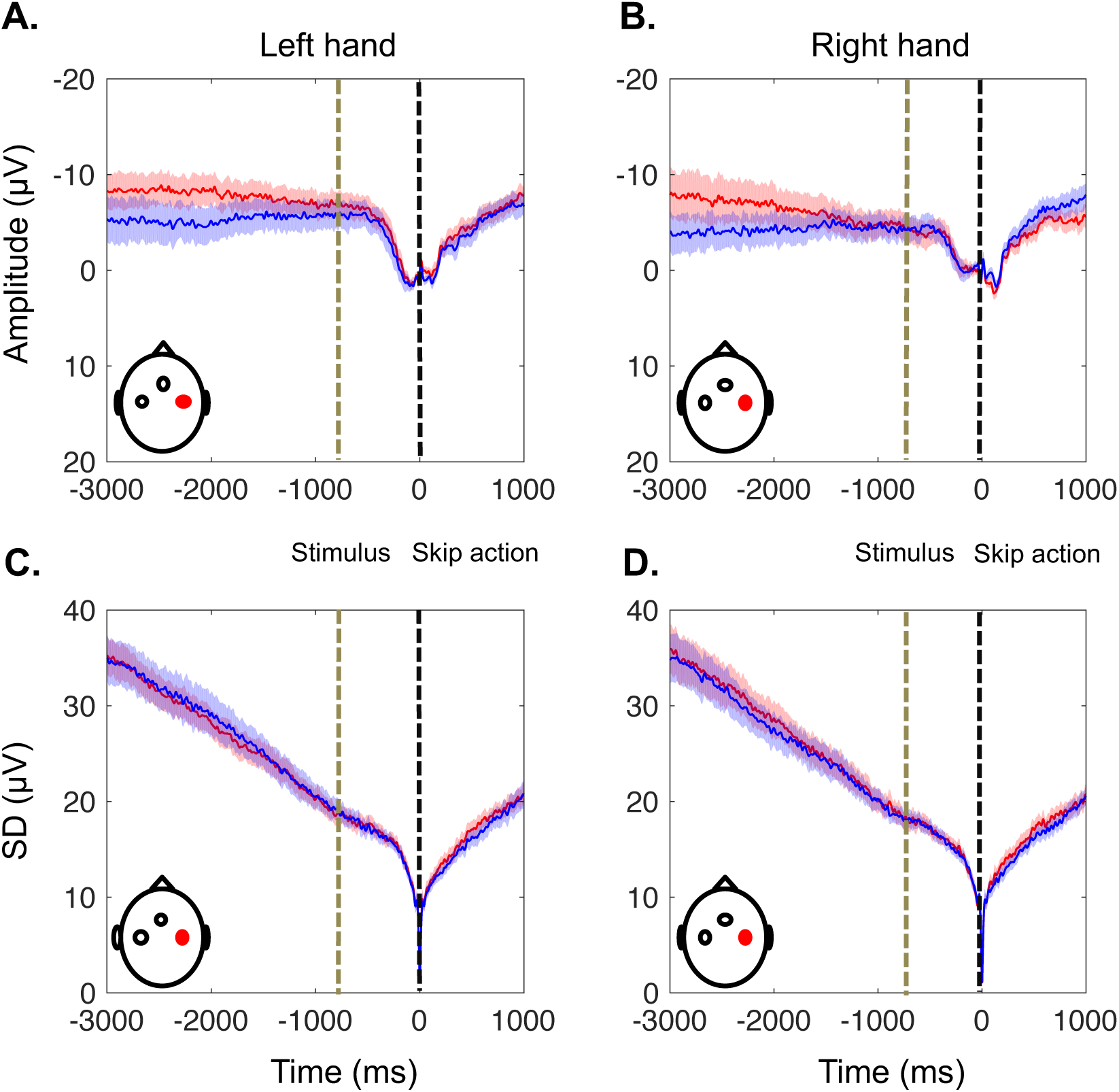
EEG activity from the effector-independent electrode (C4) prior to response to coherent dot motion. The red and blue lines represent self-initiated and externally-triggered skip conditions, respectively. Format as in Supplementary Fig. 2. A,B. Grand average RP amplitude ± standard error of the mean across participants (SEM) for the responses with the left (A) and right (B) hand. C,D. Standard deviation across trials averaged across participants ± SEM for the responses with the left (C) and right (D) hand.

**Supplementary Table 1.** Mean of waiting time before skipping with left and right hand in self-initiated and externally-triggered conditions in Exp.1.

| Subject | Mean wait (s)<br>self-initiated<br>left-hand | Mean wait (s)<br>self-initiated<br>right-hand | Mean wait (s)<br>externally-triggered<br>left-hand | Mean wait (s)<br>externally-triggered<br>right-hand |
| --- | --- | --- | --- | --- |
| 1 | 5.33 | 6.95 | 5.68 | 7.31 |
| 2 | 8.18 | 7.86 | 8.72 | 8.39 |
| 3 | 5.07 | 5.99 | 5.53 | 6.45 |
| 4 | 5.90 | 6.98 | 6.31 | 7.31 |
| 5 | 6.25 | 6.77 | 6.75 | 7.25 |
| 6 | 7.61 | 6.56 | 8.06 | 6.90 |
| 7 | 5.43 | 7.75 | 5.76 | 8.17 |
| 8 | 6.08 | 7.27 | 6.35 | 7.88 |
| 9 | 6.34 | 6.84 | 7.10 | 7.41 |
| 10 | 6.35 | 6.37 | 6.79 | 6.77 |
| 11 | 6.25 | 6.87 | 6.66 | 7.30 |
| 12 | 6.97 | 7.16 | 7.66 | 7.85 |
| 13 | 7.18 | 7.10 | 7.89 | 7.77 |
| 14 | 8.73 | 7.41 | 9.49 | 8.14 |
| 15 | 8.84 | 8.09 | 9.51 | 8.86 |
| 16 | 6.98 | 6.38 | 7.67 | 7.13 |
| 17 | 6.88 | 6.52 | 7.54 | 7.23 |
| 18 | 6.29 | 6.08 | 6.85 | 6.71 |
| 19 | 6.93 | 7.04 | 7.63 | 7.73 |
| 20 | 6.02 | 6.17 | 6.56 | 6.69 |

**Supplementary Table 2.** Mean of waiting time before skipping in Exp.2.

| Subject | Mean wait (s)<br>self-initiated<br>limited | Mean wait (s)<br>externally-triggered<br>limited | Mean wait (s)<br>self-initiated<br>unlimited | Mean wait (s)<br>externally-triggered<br>unlimited |
| --- | --- | --- | --- | --- |
| 1 | 9.60 | 10.01 | 6.60 | 7.06 |
| 2 | 12.81 | 13.30 | 8.78 | 9.18 |
| 3 | 12.42 | 13.14 | 9.71 | 10.14 |
| 4 | 8.06 | 8.57 | 5.57 | 5.98 |
| 5 | 13.11 | 13.49 | 11.33 | 11.90 |
| 6 | 8.55 | 9.37 | 6.88 | 7.24 |
| 7 | 13.77 | 14.15 | 7.09 | 7.56 |
| 8 | 12.42 | 12.65 | 7.91 | 8.22 |
| 9 | 9.05 | 9.52 | 6.35 | 7.19 |
| 10 | 7.53 | 7.86 | 4.26 | 4.56 |
| 11 | 14.59 | 15.71 | 9.05 | 9.66 |
| 12 | 11.78 | 12.09 | 7.59 | 8.42 |
| 13 | 11.62 | 11.98 | 6.41 | 6.97 |
| 14 | 10.63 | 10.49 | 5.41 | 5.94 |
| 15 | 13.68 | 14.06 | 8.72 | 9.52 |
| 16 | 7.60 | 8.38 | 6.93 | 7.45 |
| 17 | 13.30 | 13.63 | 8.07 | 9.13 |
| 18 | 11.95 | 11.88 | 7.46 | 8.02 |
| 19 | 12.22 | 13.36 | 9.62 | 9.96 |

